# Quantifying the effect of caloric and non-caloric sweeteners in the brain response using EEG and convolutional neural network

**DOI:** 10.1101/2021.10.25.465723

**Authors:** Gustavo Voltani von Atzingen, Hubert Luzdemio Arteaga Miñano, Amanda Rodrigues da Silva, Nathalia Fontanari Ortega, Ernane José Xavier Costa, Ana Carolina de Sousa Silva

## Abstract

Sweetener type can influence sensory properties and consumer’s acceptance and preference for low-calorie products. An ideal sweetener does not exist, and each sweetener must be used in situations to which it is best suited. Aspartame and sucralose can be good substitutes for sucrose in passion fruit juice. Despite the interest in artificial sweeteners, little is known about how artificial sweeteners are processed in the human brain. Here, we evaluated brain signals of 11 healthy subjects when they tasted passion fruit juice equivalently sweetened with sucrose (9.4 g/100 g), sucralose (0.01593 g/100 g), or aspartame (0.05477 g/100 g). Electroencephalograms were recorded for two sites in the gustatory cortex (i.e., C3 and C4). Data with artifacts were disregarded, and the artifact-free data were used to feed a CNN. Our results indicated that the brain responses distinguish juice sweetened with different sweeteners with an average accuracy of 0.823.

**Practical Applications:** Finding sweeteners that best fit consumer preferences evolves understanding how the gustatory cortex processes sweeteners. Ideal equivalence will occur when the brain is no longer able to distinguish stimuli that are consciously perceived. This study presents a method of signal acquisition using a single channel and an open-source processing environment. This would allow, for example, to disregard the use of a commercial electroencephalograph and expand the studies in this area and offering to food industry additional tools in the development of products sweetened with non-caloric sweeteners.

## 1 INTRODUCTION

Sugar has been the main sweetener in human diet for centuries. It represents a high percentage of the human daily energy consumption, but it offers little additional nutritional value (Bassoli & Merlini, 2003; Caballero, 2013; Zorn et al., 2014).

Given that replacing sugar with non-caloric (or low-calorie) sweeteners have become popular among consumers seeking to lose or to maintain weight. Moreover, foods with the same sweetening capacity might be perceived differently due to their caloric content (Frank et al., 2008; Hill et al., 2014). Additionally, the brain might respond distinctly to perceptually similar and identical tastes (Andersen et al., 2019).

A satisfactory method for collecting an artifact-free EEG signal has not been developed yet, but there are important results relating gustatory stimuli to evoked potential parameters in humans (Andersen et al., 2019; Hashida et al., 2005; Jacquin-Piques et al., 2016; Linforth, 2000; Mouillot et al., 2020; Ohla et al., 2012). In addition, from the food science standpoint, several aspects must be considered when it comes to taste perception, including the hypothesis that the taste perception threshold is related not only to the sensitivity of the sensory organ—in this case, the tongue—but also to a cognitive process in the brain (Huang et al., 2006; Naim et al., 2002; Okamoto & Dan, 2007).

Any study involving equi-intense sweeteners (i.e., sweeteners used in equivalent amounts) must consider how sweeteners influence sensory properties and consumer’s acceptance and preference for low-calorie products (Pinheiro et al., 2005). An ideal sweetener does not exist, so each sweetener is appropriate for specific situations (Nabors, 2002). Passion fruit is a popular tropical fruit that has an important commercial variety named yellow passion fruit, which is used to prepare juice that requires sweetening (Deliza et al., 2005). Aspartame and sucralose can be good substitutes for sucrose in passion fruit juice (Rocha & Bolini, 2015a, 2015b).

For this study, we have hypothesized that we can distinguish the stimuli resulting from the consumption of drinks sweetened with caloric or non-caloric sweeteners in EEG signal. To test this hypothesis, we have compared brain signals acquired in response to the consumption of passion fruit juice sweetened with sucrose (caloric sweetener), sucralose, or aspartame (low-calorie sweeteners).

## 2 BRIEF REVIEW OF RELEVANT METHODS

### 2.1 Convolutional Neural Networks

Methods based on different categories of learning, including supervised, semi-supervised, and unsupervised learning, have been proposed. Supervised learning is a learning technique that uses labeled data. One of the most recent and successful techniques within supervised learning is Deep Learning (DL), a new field of machine learning that allows computational methods composed of multiple processing layers to be employed (Russakovsky et al., 2015). Convolutional Neural Networks (CNNs) are another supervised learning approach. Other supervised deep leaning approaches include Deep Neural Networks (DNNs), Recurrent Neural Networks (RNNs), like Long Short Term Memory (LSTM), and Gated Recurrent Units (GRUs) (Alom et al., 2019). The approaches differ mainly in terms of connection topologies and forms of activation.

CNNs were first proposed by Fukushima in 1988 (Fukushima, 1988), but this approach was not widely employed because computation hardware was limited for training the networks. Deep CNNs are typical feedforward neural networks to which Backpropagation (BP) algorithms are applied to adjust the network parameters (weights and biases), to reduce the cost function value (X. Li et al., 2016).

Two types of layers exist in the network low- and middle-levels: convolutional layers and max-pooling layers. The even-numbered layers are used for convolutions, whilst the odd-numbered layers are employed for max-pooling operations. The output nodes of the convolution and max-pooling layers are grouped into a 2D plane called feature mapping. Each plane of a layer is usually derived from the combination of one or more planes of previous layers. The nodes of a plane are connected to a small region of each connected plane of the previous layer. Each node of the convolution layer extracts the features from the input by convolution operations on the input nodes. As the features propagate to the highest layer or level, the dimensions of features are reduced depending on the kernel size for the convolutional and max-pooling operations, respectively. The output of the last CNN layer is used as the input for a fully connected network, which is called classification layer. In the classification layer, the extracted features are taken as inputs with respect to the dimension of the weight matrix of the final neural network (Alom et al., 2019).

Learning is achieved by adjusting the synaptic weights and by using the values from previous layers and a mathematical activation function applied on them. These hidden layers allow nonlinear and complex problems to be handled (Saeed et al., 2019). The CNN is trained with the popular BP algorithm with Stochastic Gradient Descent (SGD) (Alom et al., 2019).

Compared to traditional machine learning methods, deep learning does not require too much feature engineering because it can automatically extract features and complete tasks that can only be achieved by feature engineering in traditional machine learning algorithms (J. Li et al., 2020). This allows data representations with multiple levels of abstraction to be learned, which can dramatically improve state-of-the-art speech recognition, visual object recognition, object detection, and many other domains such as drug discovery, genomics (LeCun et al., 2015), and food engineering (Pfisterer et al., 2018). Furthermore, deep learning promotes the development of intelligent fields like image processing (Russakovsky et al., 2015), speech recognition (Collobert et al., 2011), and two-player games (Silver et al., 2017). In this study field, DNNs are an important and promising application (J. Li et al., 2020). Prominent areas include self-driving cars (Badue et al., 2021) and consumer behavior (Kim et al., 2020).

## 3 MATERIAL AND METHODS

The methods used herein were divided into three main sections: (1) Participant selection, (2) acquisition of EEG signal from the selected group, and (3) signal processing and CNN (Convolutional Neural Network) training and tests. Our aim was to use a CNN to determine differences between stimuli.

### 3.1 Stimuli

The methods used herein were divided into three main sections: (1) Participant selection, (2) acquisition of EEG signal from the selected group, and (3) signal processing and CNN (Convolutional Neural Network) training and tests. Our aim was to use a CNN to determine differences between stimuli.

### 3.2 Participant Selection

A total of 105 volunteers were included in this study. The volunteers comprised students, teachers, and employees aged 19–55 years, recruited on campus. They did not have diabetes, smoke, or use medications that affect taste or cognitive processes. Preference was given to volunteers that were used to consuming passion fruit juice (or that at least had no aversion to the fruit taste).

Each volunteer received samples containing 30 mL of passion fruit juice sweetened with different amounts of sugar (i.e., 4.7, 7.05, 9.4, 11.75, and 14.1 g). The samples were placed in disposable cups and numbered randomly between 0000 and 9999 (e.g., A7932). The participants received samples in a randomized sequence of concentrations and had to answer the following question: “How much sugar did you have in your juice?” Answers were collected in a 9.0-cm visual scale (VAS). VASs are input mechanisms that allow users to specify a value within a predefined range. The volunteers were instructed to consider the centre of the scale as ideal sweetness, 0 as less sweet than ideal, and 9 as sweeter than ideal. Our objective was to select individuals with a perception of sugar ideal sweetness as close as possible to 9.4 g/100 g and, among these individuals, to select those that had good ability to order the samples according to the sugar concentration.

The volunteers were informed about the nature and aims of the experiments and provided informed consent. The study was approved by the Ethics Committee of the Animal Science and Food Engineering College (FZEA) of the University of São Paulo (USP) (CAAE 59017516.6.0000.5422).

#### 3.2.1 EEG recording

EEG signals were acquired from 11 individuals (both sexes) while they were tasting passion fruit juice. The individuals had been selected according to item ‘Participant selection’ and had signed an informed consent form (ICF).

EEG was carried out non-invasively on the scalp surface; the participants wore an EEG cap. The signal was acquired at positions C3 and C4 in the primary gustatory cortex (Hashida et al., 2005; Kobayakawa, 1999) as defined by the international 10–20 electrode disposal system (Jasper and H., 1958). This electrode position was chosen on the basis of the study by Kabayakawa et al. (1999), who showed magnetic fields recorded from the brain (i.e., MEG) in response to two tastants: 1 M NaCl and 3 mM saccharin, one of which was more activated in the central area of the head at positions C3 and C4. The ground electrodes were placed on the participant’s ear lobes, and a reference electrode was placed on the forehead. The signals were sampled at 512 Hz by using an iCelera digital portable electroencephalograph (iBlue 52). Each recording lasted 16 seconds.

The volunteers were accommodated inside a Faraday cage, and the EEG signal started being recorded when the volunteer drank the solution (juice). The volunteers were instructed to keep their eyes closed and not to move during the recordings. Each volunteer received eight 30-mL samples for evaluation. The samples were placed in disposable cups and numbered randomly between 0000 and 9999 (e.g., A7932). The samples were offered in duplicate in the following order: water (reference), passion fruit juice sweetened with sucrose, passion fruit juice sweetened with sucralose, and passion fruit juice sweetened with aspartame. Each volunteer participated on three different days of the experiment (repetitions). In the intervals between the sweetened samples, the participants received sparkling water to clean residues from their palate and to reduce other interferences.

#### 3.2.2 Signal processing and CNN training

The signal was processed by using the Python programming language and the Pandas, NumPy, and SciPy libraries. These libraries are employed to solve mathematical and scientific problems, including signal processing. Initially, the data were inspected, and the recordings with many artifacts were excluded from the database. The remaining data consisted of signals from 68 experiments lasting 16 seconds each, recorded in two channels (i.e., C3 and C4). These signals came from one of four stimuli (i.e., sucrose, sucralose, aspartame, or water) and were captured across three days of repetitions per volunteer. The data were bandpass-filtered from 8 to 40 Hz, and each signal was divided into two-second segments with a 0.1 second stride. This means that 16-second segments (512 samples per second) were resampled in 9520 vectors at a length of 1028. The EEG dataset and the processing code that supported the findings of this study are available on GitHub (https://github.com/Atzingen/EEG_Sweetners.).

When the vectors with length *1028* were used as inputs for the CNN, they were structured as a network consisting of three parallel convolution processes merged into a fully two-layer-deep connected network.

The deep learning model was built by using an open-source Python library, i.e., Keras, which runs in a TensorFlow background. Optimal performance (e.g., number of neurons on a given layer and number of filters and kernel size on the convolution layer) was determined by using manual fine adjustment of the hyperparameters (e.g., number of neurons per fully connected layer and number of filters and kernel size for convolutional layers).

The last network layer contained four neurons, which were responsible for mapping four possible results of an entry (water, sugar, aspartame, or sucralose). A categorical cross-entropy loss function was used.

Training was accomplished by using an Adam algorithm with two-thirds of the data, while the test was performed with the remaining one-third.

Finally, performance evaluation metrics, such as accuracy, precision, recall and F1 score were calculated as follows.

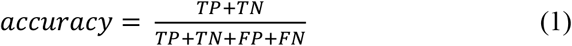

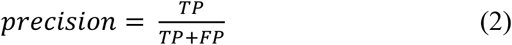

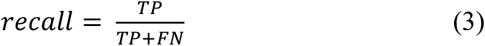

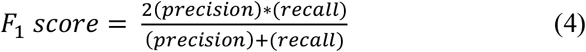

where TP (true positive), FP (false positive), TN (true negative) and FN (false negative) are technical terms for binary classifier. Specific, TP is the positive samples correctly classified, FP is the positive samples misclassified, TN is the negative samples correctly classified and FN represents the negative samples misclassified.

## 4 RESULTS AND DISCUSSION

### 4.1 Participant selection

We selected the groups according to two criteria, namely, preference for the sample with a sucrose concentration of 9.4 g/100 g and good ability to order the sample concentrations. This means that the selected participants indicated values around 4.5 cm on the VAS, which was equivalent to a sucrose concentration of 9.4 g/100 g. Furthermore, they were able to order all the concentrations properly. Of the individuals considered fit, 11 agreed to participate in the brain signal acquisition stage.

Figure 1-a illustrates the scale presented to the participants. Figure 1-b shows participant A, who placed all the samples in the correct sequence of concentrations and chose the sample with sugar concentration of 9.4 g/100 g as his favourite. He was selected for the next step. Participants B and C were not selected. Participant B (Figure 1-c) placed the samples correctly, but his preferred sample was not the sample with sugar concentration of 9.4 g/100 g. Although participant C (Figure 1-d) preferred the sample with a sugar concentration of 9.4 g/100 g, he was not able to order the samples correctly.

**Figure 1:**
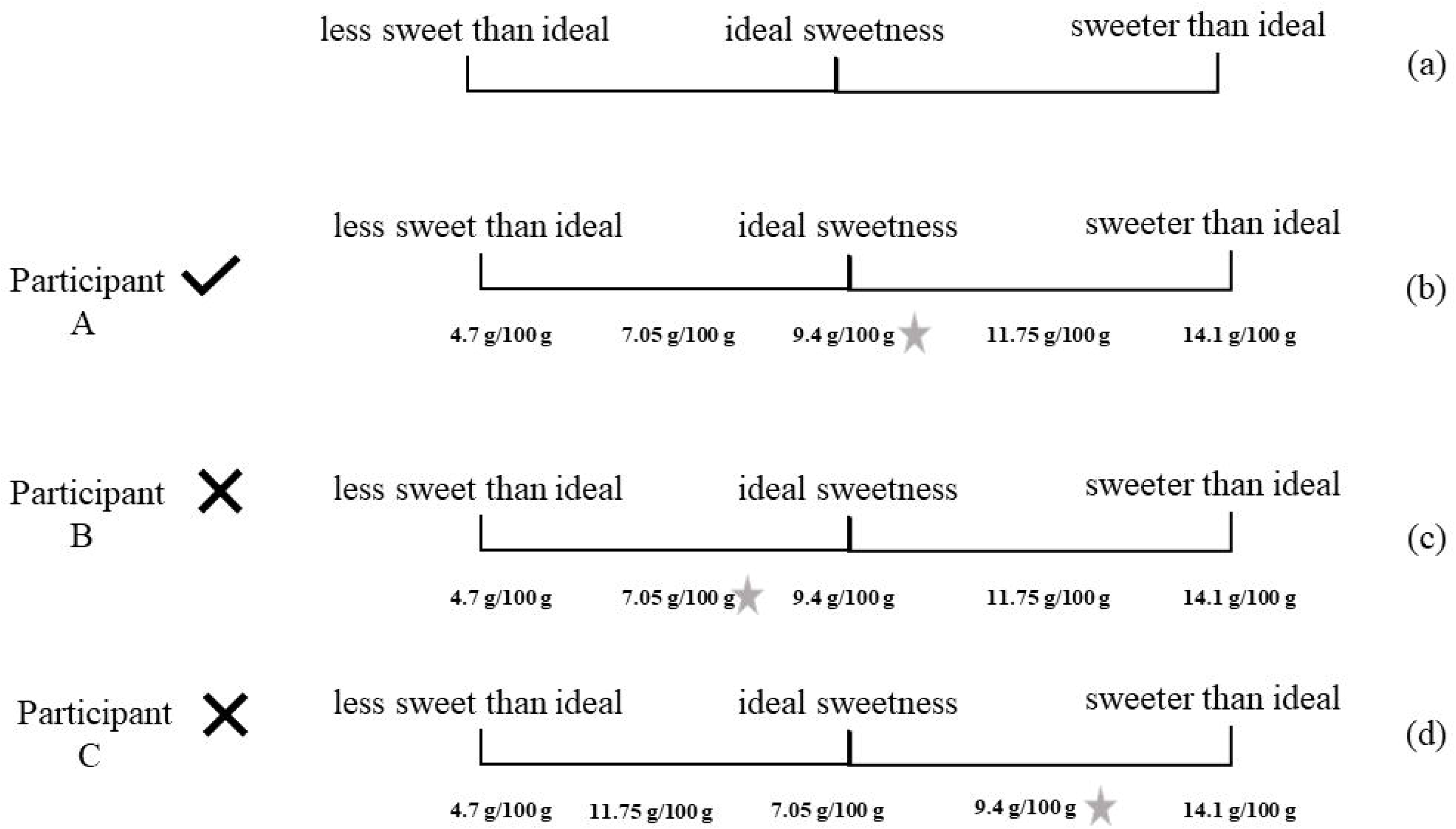
(a) Participant selection form. (b) Participant selection form from a selected participant. (c) Participant selection form from a discarded participant. (d) Participant selection form from a discarded participant. The star placed on the right side illustrates the preferred sample. In the original form, the volunteer marked the sample number on the scale. For didactic purposes, these codes were replaced with the sample concentration in the figure.

### 4.2 Signal processing and CNN training

Figure 2 shows the network architecture that achieved the best performance. The best number for *n* was 20, as can be seen by the input layer shape. The other three fully connected layers had 16, 16, and 64 neurons. Its dropout optimal value was 0.2.

**Figure 2:**
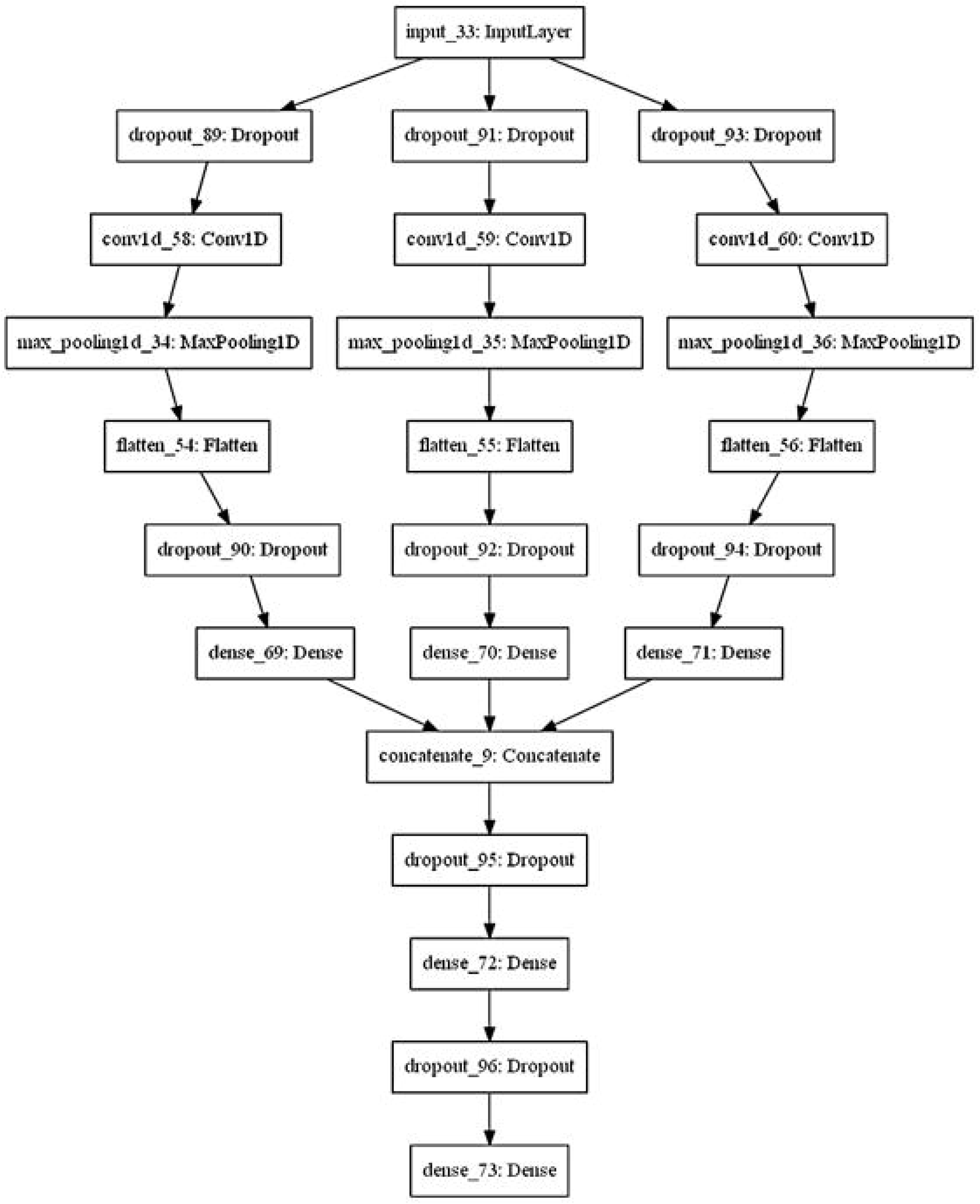
The convolutional neural network (CNN) architecture that achieved the best performance.

We used this optimal architecture to train the CNN with 70 % of the data, while we employed the remaining 30% for the test.

For visualization, Figure 3 presents the confusion matrix for this dataset. In the confusion matrix, the horizontal axis is the predicted label, and the vertical axis is the true label. The elements on the diagonal represent the numbers of correctly classified samples.

**Figure 3:**
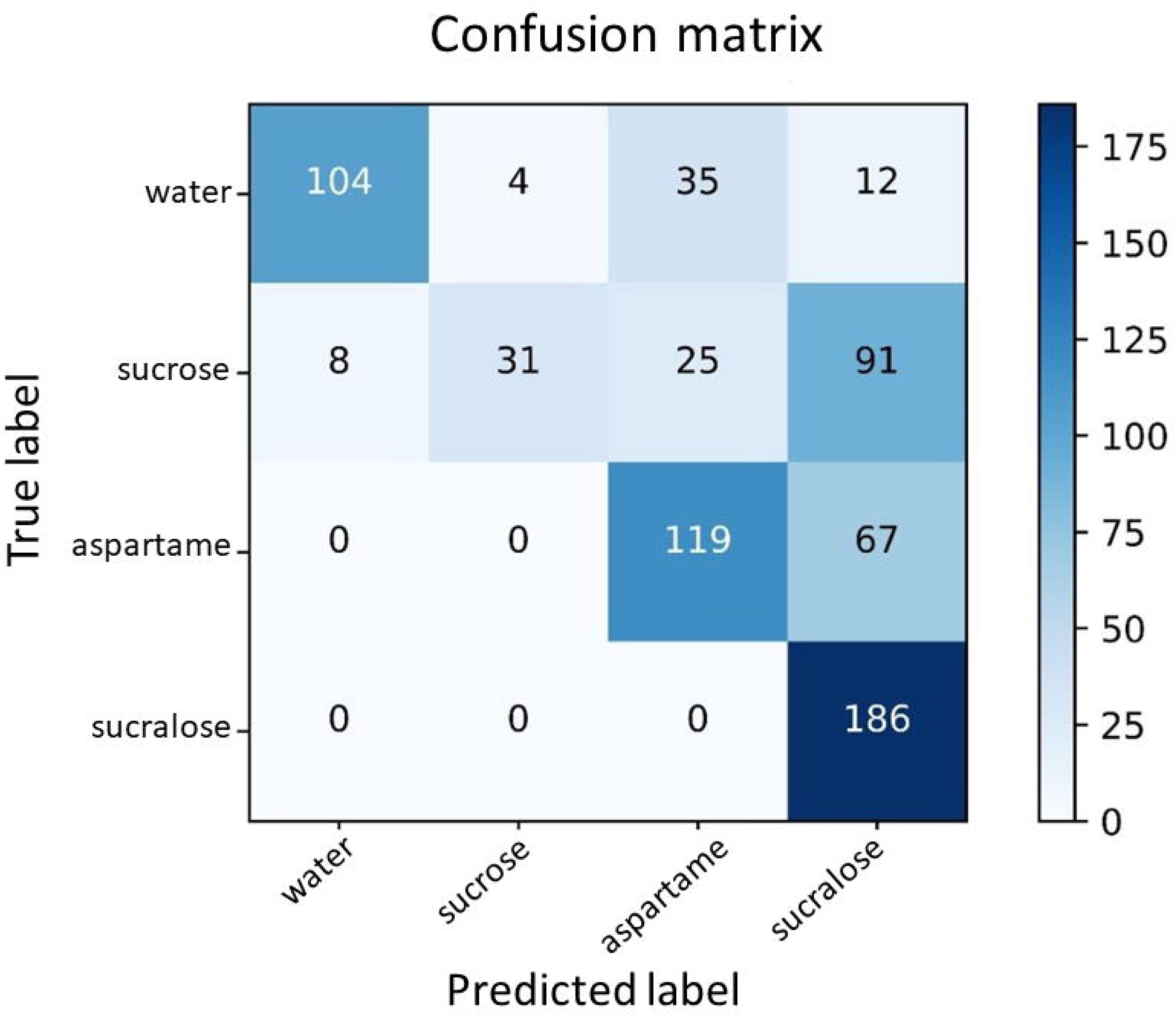
Confusion matrix for the classes water, sucrose, sucralose, and aspartame.

In Figure 3, for the water class, 104 samples were correctly predicted as water (67.1%), whereas 35 samples (7.7%) were incorrectly predicted as aspartame and 4 samples (2.6%) were incorrectly predicted as sucralose. For the sucrose class, 31 samples (20.0%) were correctly predicted as sucrose, whilst 25 samples (16.1%) were incorrectly predicted as aspartame, and 91 samples (58.7%) were incorrectly predicted as sucralose. As for the aspartame class, 119 samples (64.0%) were correctly classified, while 67 samples (36.0%) were incorrectly predicted as sucralose. Concerning the sucralose class, 186 samples (100.0%) were correctly classified. The main difficulty lay in the classes of water and sucrose.

Table 1 lists the metrics results for the four stimuli (classes).

**Table 1.** DNN hits for the pairs of taste stimuli.

|  | **Water** | **Sucrose** | **Sucralose** | **Aspartame** |
| --- | --- | --- | --- | --- |
| **Water** | - | 70.71 | 58.92 | 60.53 |
| **Sucrose** | 70.71 | - | 63.23 | 65.95 |
| **Sucralose** | 58.92 | 63.23 | - | 65.98 |
| **Aspartame** | 60.53 | 65.95 | 65.28 | - |

The average metrics (Table 1) were: 0.823 for accuracy, 0.750 for precision, 0.628 for recall and 0.611 for F_1_ score. There are no studies using CNN in similar databases for comparison purposes, but these values are compatible with those obtained for studies involving brain signals to assess emoticons (Yin et al., 2017) and cerebral dominance (Toraman et al., 2019).

**Table 1:** Results of CNN classification performance for the four stimuli (classes).

|  | Accuracy | Precision | Recall | F<sub>1</sub> score |
| --- | --- | --- | --- | --- |
| Water | 0.9135 | 0.9286 | 0.6710 | 0.7790 |
| Sucrose | 0.8123 | 0.8857 | 0.2000 | 0.3263 |
| Aspartame | 0.8138 | 0.6648 | 0.6398 | 0.6521 |
| Sucralose | 0.7507 | 0.5225 | 1.0000 | 0.6863 |

When we consider reference (water) only, the classification in Figure 3 indicated that the average identification accuracy is 91,4% compared to an overall classification of 82.3%. This work uses a drink instead of a solution and, in a first study of this nature, it was important to be able to distinguish a critical reference well.

When we compare general performance with reference performance, mainly f_1_ score that consider recall and precision measures simultaneously we can deduce that misclassifications in sweeteners classes are greater than that in reference class.

Analysing Figure 3 in more detail, suggests that the network is more sensitive to the low-calorie sweetener classes than to the sucrose class. By evaluating the false positives (FP) of the classification, aspartame can be predicted to be sucralose, but not sugar. In turn, sucralose had no false positives. The most surprising result is that sucrose can be classified both as sugar and low-calorie sweetener. In other words, evaluated low-calorie sweeteners had not been confused with the caloric sweetener, but the caloric sweetener can be confused with the low-calorie sweetener. Andersen et al. (2019) observed that similar tastes that are consciously indistinguishable can result in different brain cortical activations. A similar result was obtained when gustatory evoked potentials (GEPs) were used to assess the brain response to sucrose, aspartame, and stevia in humans (Mouillot et al., 2020). The authors stated that, although sucrose, aspartame, and stevia led to the same taste perception, the GEPs showed that cerebral activation by these different sweet solutions had different recordings. They suggested that, besides the difference in taste receptors and cerebral areas activated by these substances, neural plasticity and changes in the synaptic connections related to sweet innate preference and sweet conditioning could explain the differences in cerebral gustatory processing after sucrose and sweetener activation. The results presented herein showed that this may be true.

Unlike Andersen et al. (2019) and Mouillot et al. (2020), in this study we evaluated a beverage (i.e., passion fruit juice) with good acceptance after it is sweetened with aspartame or sucralose (Rocha & Bolini, 2015a, 2015b) as a stimulus instead of evaluating a sweet solution. This may explain FP rates in classification. During tasting, the stimuli came not only from the sweet solution, but also from the other sensory characteristics of the beverage.

Apart from using a beverage instead of a solution and from dispensing a taste delivery system (Andersen et al., 2019; Frank et al., 2008; Jacquin-Piques et al., 2016; Kobayakawa et al., 1996; Kobayakawa, 1999) that is common in this kind of experimental design, we only placed two electrodes near the gustatory cortex. While this is not a new strategy (Cincotti et al., 2002; Costa & Cabral, 2000; Hashida et al., 2005), it is uncommon (Andersen et al., 2019; Frank et al., 2008; Jacquin-Piques et al., 2016; Kobayakawa et al., 1996; Kobayakawa, 1999) and has advantages and disadvantages. The main disadvantage is that it reduces the amount of data and makes some feature extraction techniques unfeasible. On the other hand, developing commercial applications based on a single EEG channel, such as the applications used by Hashida et al. (2005) and Silva et al. (2005), is easy.

Andersen et al. (2019) observed that response discrimination by between‐participant qEEG is inferior to discrimination by within‐participant qEEG. Therefore, we can improve the results of this study by increasing the number of samples tested per individual and also the number of individuals. This would allow us to train the network for each participant, for instance. However, this approach would involve the use of a taste delivery system to facilitate the increase in the number of samples.

This fact does not invalidate the main contribution of this study, which corroborates recent studies (Andersen et al., 2019; Frank et al., 2008; Mouillot et al., 2020) stating that similar stimuli, despite being consciously indistinguishable, may result in different cortical responses. Moreover, this study included a beverage instead of a solution and used only two monitoring positions (i.e., C3 and C4) over the gustatory cortex.

## 5 CONCLUSION

We have compared brain signals acquired in response to the consumption of passion fruit juice sweetened with sucrose (caloric sweetener), sucralose, or aspartame (low-calorie sweeteners). We used the artifact-free data to feed a CNN. The results indicated that the brain responses can distinguish the juice sweetened with sucrose from the juice sweetened with aspartame and sucralose.

## Acknowledgments

This work was supported by Fundação de Amparo à Pesquisa do Estado de São Paulo - FAPESP under the grant number 2018/03027-0.

We also thank Ajinomoto for the donation of aspartame (AminoSweet^TM^).

## REFERENCES

Alom, M. Z., Taha, T. M., Yakopcic, C., Westberg, S., Sidike, P., Nasrin, M. S., Hasan, M., Van Essen, B. C., Awwal, A. A. S., & Asari, V. K. (2019). A State-of-the-Art Survey on Deep Learning Theory and Architectures. Electronics, 8(3), 292. https://doi.org/10.3390/electronics8030292

Andersen, C. A., Kring, M. L., Andersen, R. H., Larsen, O. N., Kjaer, T. W., Kidmose, U., Møller, S., & Kidmose, P. (2019). EEG discrimination of perceptually similar tastes. Journal of Neuroscience Research, 97(3), 241–252. https://doi.org/10.1002/jnr.24281

Badue, C., Guidolini, R., Carneiro, R. V., Azevedo, P., Cardoso, V. B., Forechi, A., Jesus, L., Berriel, R., Paixão, T. M., Mutz, F., de Paula Veronese, L., Oliveira-Santos, T., & De Souza, A. F. (2021). Self-driving cars: A survey. In Expert Systems with Applications (Vol. 165, p. 113816). Elsevier Ltd. https://doi.org/10.1016/j.eswa.2020.113816

Bassoli, A., & Merlini, L. (2003). SWEETENERS | Intensive. In Encyclopedia of Food Sciences and Nutrition (pp. 5688–5695). Elsevier. https://doi.org/10.1016/B0-12-227055-X/01172-X

Caballero, B. (2013). Sucrose: Dietary Sucrose and Disease. In Encyclopedia of Human Nutrition (pp. 231–233). Elsevier. https://doi.org/10.1016/B978-0-12-375083-9.00257-9

Cincotti, F., Mattia, D., Babiloni, C., Carducci, F., Bianchi, L., del R Millán, J., Mouriño, J., Salinari, S., Marciani, M. G., & Babiloni, F. (2002). Classification of EEG mental patterns by using two scalp electrodes and Mahalanobis distance-based classifiers. Methods of Information in Medicine, 41(4), 337–341. http://www.ncbi.nlm.nih.gov/pubmed/12425246

Collobert, R., Weston, J., Bottou, L., Karlen, M., Kavukcuoglu, K., & Kuksa, P. (2011). Natural language processing (almost) from scratch. Journal of Machine Learning Research, 12, 2493–2537.

Costa, E. J. X., & Cabral, E. F. (2000). EEG-based discrimination between imagination of left and right hand movements using adaptive gaussian representation. Medical Engineering & Physics, 22(5), 345–348. https://doi.org/10.1016/S1350-4533(00)00051-5

Deliza, R., Macfie, H., & Hedderley, D. (2005). THE CONSUMER SENSORY PERCEPTION OF PASSION-FRUIT JUICE USING FREE-CHOICE PROFILING. Journal of Sensory Studies, 20(1), 17–27. https://doi.org/10.1111/j.1745-459X.2005.050604.x

Frank, G. K. W., Oberndorfer, T. A., Simmons, A. N., Paulus, M. P., Fudge, J. L., Yang, T. T., & Kaye, W. H. (2008). Sucrose activates human taste pathways differently from artificial sweetener. NeuroImage, 39(4), 1559–1569. https://doi.org/10.1016/j.neuroimage.2007.10.061

Fukushima, K. (1988). Neocognitron: A hierarchical neural network capable of visual pattern recognition. Neural Networks, 1(2), 119–130. https://doi.org/10.1016/0893-6080(88)90014-7

Hashida, J. C., Silva, A. C. de S., Souto, S., & Costa, E. J. X. (2005). EEG pattern discrimination between salty and sweet taste using adaptive Gabor transform. Neurocomputing, 68, 251–257. https://doi.org/10.1016/J.NEUCOM.2005.04.004

Hill, S. E., Prokosch, M. L., Morin, A., & Rodeheffer, C. D. (2014). The effect of non-caloric sweeteners on cognition, choice, and post-consumption satisfaction. Appetite, 83, 82–88. https://doi.org/10.1016/j.appet.2014.08.003

Huang, A. L., Chen, X., Hoon, M. A., Chandrashekar, J., Guo, W., Tränkner, D., Ryba, N. J. P., & Zuker, C. S. (2006). The cells and logic for mammalian sour taste detection. Nature, 442(7105), 934–938. https://doi.org/10.1038/nature05084

Jacquin-Piques, A., Gaudillat, S., Mouillot, T., Gigot, V., Meillon, S., Leloup, C., Penicaud, L., & Brondel, L. (2016). Prandial States Modify the Reactivity of the Gustatory Cortex Using Gustatory Evoked Potentials in Humans. Frontiers in Neuroscience, 9. https://doi.org/10.3389/fnins.2015.00490

Jasper, & H., H. (1958). The ten twenty electrode system of the international federation. Electroencephalography and Clinical Neurophysiology, 10, 371–375. http://ci.nii.ac.jp/naid/10020218106/en/

Kim, J., Ji, H. G., Oh, S., Hwang, S., Park, E., & del Pobil, A. P. (2020). A deep hybrid learning model for customer repurchase behavior. Journal of Retailing and Consumer Services, 102381. https://doi.org/10.1016/j.jretconser.2020.102381

Kobayakawa, T. (1999). Spatio-temporal Analysis of Cortical Activity Evoked by Gustatory Stimulation in Humans. Chemical Senses, 24(2), 201–209. https://doi.org/10.1093/chemse/24.2.201

Kobayakawa, T., Endo, H., Ayabe-Kanamura, S., Kumagai, T., Yamaguchi, Y., Kikuchi, Y., Takeda, T., Saito, S., & Ogawa, H. (1996). The primary gustatory area in human cerebral cortex studied by magnetoencephalography. Neuroscience Letters, 212(3), 155–158. https://doi.org/10.1016/0304-3940(96)12798-1

LeCun, Y., Bengio, Y., & Hinton, G. (2015). Deep learning. Nature, 521(7553), 436–444. https://doi.org/10.1038/nature14539

Li, J., Ping, Y., Li, H., Li, H., Liu, Y., Liu, B., & Wang, Y. (2020). Prognostic prediction of carcinoma by a differential-regulatory-network-embedded deep neural network. Computational Biology and Chemistry, 107317. https://doi.org/10.1016/j.compbiolchem.2020.107317

Li, X., Zhang, G., Li, K., & Zheng, W. (2016). Deep Learning and Its Parallelization. In Big Data (pp. 95–118). Elsevier. https://doi.org/10.1016/B978-0-12-805394-2.00004-0

Linforth, R. S. (2000). Developments in instrumental techniques for food flavour evaluation: future prospects. Journal of the Science of Food and Agriculture, 80(14), 2044–2048. https://doi.org/10.1002/1097-0010(200011)80:14<2044::AID-JSFA753>3.0.CO;2-Z

Mouillot, T., Parise, A., Greco, C., Barthet, S., Brindisi, M.-C., Penicaud, L., Leloup, C., Brondel, L., & Jacquin-Piques, A. (2020). Differential Cerebral Gustatory Responses to Sucrose, Aspartame, and Stevia Using Gustatory Evoked Potentials in Humans. Nutrients, 12(2), 322. https://doi.org/10.3390/nu12020322

Nabors, L. O. (2002). Sweet Choices: Sugar Replacements for Foods and Beverages. Food Technology, 56(7), 28–35. https://www.ift.org/news-and-publications/food-technology-magazine/issues/2002/july/features/sweet-choices-sugar-replacements-for-foods-and-beverages

Naim, M., Nir, S., Spielman, A. I., Noble, A. C., Peri, I., Rodin, S., & Samuelov-Zubare, M. (2002). Hypothesis of Receptor-Dependent and Receptor-Independent Mechanisms for Bitter and Sweet Taste Transduction: Implications for Slow Taste Onset and Lingering Aftertaste (pp. 2–17). https://doi.org/10.1021/bk-2002-0825.ch001

Ohla, K., Busch, N. A., & Lundström, J. N. (2012). Time for Taste—A Review of the Early Cerebral Processing of Gustatory Perception. Chemosensory Perception, 5(1), 87–99. https://doi.org/10.1007/s12078-011-9106-4

Okamoto, M., & Dan, I. (2007). Functional near-infrared spectroscopy for human brain mapping of taste-related cognitive functions. Journal of Bioscience and Bioengineering, 103(3), 207–215. https://doi.org/10.1263/jbb.103.207

Pfisterer, K. J., Amelard, R., Chung, A. G., & Wong, A. (2018). A new take on measuring relative nutritional density: The feasibility of using a deep neural network to assess commercially-prepared puréed food concentrations. Journal of Food Engineering, 223, 220–235. https://doi.org/10.1016/j.jfoodeng.2017.10.016

Pinheiro, M. V. S., Oliveira, M. N., Penna, A. L. B., & Tamime, A. Y. (2005). The effect of different sweeteners in low-calorie yogurts - a review. International Journal of Dairy Technology, 58(4), 193–199. https://doi.org/10.1111/j.1471-0307.2005.00228.x

Rocha, I. F. de O., & Bolini, H. M. A. (2015a). Passion fruit juice with different sweeteners: sensory profile by descriptive analysis and acceptance. Food Science & Nutrition, 3(2), 129–139. https://doi.org/10.1002/fsn3.195

Rocha, I. F. de O., & Bolini, H. M. A. (2015b). Different sweeteners in passion fruit juice: Ideal and equivalent sweetness. LWT - Food Science and Technology, 62(1), 861–867. https://doi.org/10.1016/J.LWT.2014.10.055

Russakovsky, O., Deng, J., Su, H., Krause, J., Satheesh, S., Ma, S., Huang, Z., Karpathy, A., Khosla, A., Bernstein, M., Berg, A. C., & Fei-Fei, L. (2015). ImageNet Large Scale Visual Recognition Challenge. International Journal of Computer Vision, 115(3), 211–252. https://doi.org/10.1007/s11263-015-0816-y

Saeed, S., Naz, S., & Razzak, M. I. (2019). An Application of Deep Learning in Character Recognition: An Overview (pp. 53–81). https://doi.org/10.1007/978-3-030-11479-4_3

Silva, A. C. de S., Arce, A. I. C., Souto, S., & Costa, E. J. X. (2005). A wireless floating base sensor network for physiological responses of livestock. Computers and Electronics in Agriculture, 49(2), 246–254. https://doi.org/10.1016/J.COMPAG.2005.05.004

Silver, D., Schrittwieser, J., Simonyan, K., Antonoglou, I., Huang, A., Guez, A., Hubert, T., Baker, L., Lai, M., Bolton, A., Chen, Y., Lillicrap, T., Hui, F., Sifre, L., van den Driessche, G., Graepel, T., & Hassabis, D. (2017). Mastering the game of Go without human knowledge. Nature, 550(7676), 354–359. https://doi.org/10.1038/nature24270

Toraman, S., Tuncer, S. A., & Balgetir, F. (2019). Is it possible to detect cerebral dominance via EEG signals by using deep learning? Medical Hypotheses, 131. https://doi.org/10.1016/J.MEHY.2019.109315

Yin, Z., Zhao, M., Wang, Y., Yang, J., & Zhang, J. (2017). Recognition of emotions using multimodal physiological signals and an ensemble deep learning model. Computer Methods and Programs in Biomedicine, 140, 93–110. https://doi.org/10.1016/J.CMPB.2016.12.005

Zorn, S., Alcaire, F., Vidal, L., Giménez, A., & Ares, G. (2014). Application of multiple-sip temporal dominance of sensations to the evaluation of sweeteners. Food Quality and Preference, 36, 135–143. https://doi.org/10.1016/j.foodqual.2014.04.003

